# Complex Properties of Training Stimuli Affect Brain Alignment in a Deep Network Model of Mouse Visual Cortex

**DOI:** 10.1101/2024.10.24.620142

**Authors:** Parsa Torabian, Yinghan Chen, Candace Ng, Stefan Mihalas, Michael Buice, Shahab Bakhtiari, Bryan Tripp

## Abstract

Deep convolutional neural networks are important models of the visual cortex that account relatively well for brain activity and are able to perform ethologically relevant functions. However, it is unknown which combination of factors, such as network architecture, training objectives, and data best align this family of models with the brain. Here we investigate the statistics of training data. We hypothesized that stimuli that are naturalistic for mice would lead to higher similarity between deep network models and activity in mouse visual cortex. We used a video-game engine to create training datasets in which we varied the naturalism of the environment, the movement statistics, and the optics of the modelled eye. The naturalistic environment substantially and consistently led to greater brain similarity, while the other factors had more subtle and area-specific effects. We then hypothesized that differences in brain similarity between the two environments arose due to differences in spatial frequency spectra, distributions of color and orientation, and/or temporal autocorrelations. To test this, we created abstract environments, composed of cubes and spheres, that resembled the naturalistic and non-naturalistic environments in these respects. Contrary to our expectations, these factors accounted poorly for differences in brain similarity due to the naturalistic and non-naturalistic environments. This suggests that the higher brain similarities we observed after training with the naturalistic environment were due to more complex factors.

## 1 Introduction

Deep convolutional neural networks have structural parallels with the visual cortex, and their internal activity has so far provided the most accurate predictions of visual cortex activity (Yamins and DiCarlo, 2016; Shi et al., 2019; Schrimpf et al., 2020). As a step in developing accurate models of brain function, it is important to understand the network configuration factors that improve these predictions. Two important factors are network structure (Schrimpf et al., 2020; Kubilius et al., 2019) and training objective (Zhuang et al., 2021; Lindsay et al., 2022a; Nayebi et al., 2023). Deep networks have also been regularized to better reflect certain properties of brain activity (Gerum et al., 2022), and network mechanisms grounded in neuroscience have been incorporated to make certain response properties more realistic (Dutta et al., 2018; Khan et al., 2023; Lindsay et al., 2022b; Schrimpf et al., 2024).

Statistics of the training stimuli may also affect similarity with brain activity. For example, training with moving stimuli and a suitable objective function can lead to the development of a model that better aligns with motion processing areas of the brain (Mineault et al., 2021), and deep networks’ activity is better aligned with mouse visual cortex when their inputs have realistic resolution for mice rather than humans (Nayebi et al., 2023). We hypothesized that training data with other naturalistic properties would result in more mouse brain-like activity.

In this study, we used MouseNet (Shi et al., 2022), a biologically grounded network architecture. To train the model, we used a predictive self-supervised learning approach (Han et al., 2019) that has been shown effective in modeling mouse visual cortex (Bakhtiari et al., 2021). We varied several stimulus properties and investigated their effect on model-brain alignment. Specifically, we created egocentric video datasets with naturalistic (for mice) vs. non-naturalistic environments, movement statistics, and optics of the eye model. These differences significantly affected the model’s alignment with mouse brain activity. The environment had the greatest and most consistent im-pact of these factors. The highest brain similarity was obtained by combining all three naturalistic conditions: environment, optics, and motion. Naturalistic motion impaired brain similarity on average, but improved similarity with the area VISam, which is particularly known to be motion selective (Marshel et al., 2011; Sit and Goard, 2020).

It is unclear which differences between the naturalistic and non-naturalistic environments account for the resulting differences in brain similarity. We suspected that these differences could be explained in terms of intuitive statistical properties, including spatial frequency spectra, colour distributions, orientation distributions, and temporal autocorrelations. However, using more abstract virtual environments, we found that this combination of properties did not account for environment-related differences in brain similarity.

## 2 Methods

Figure 1 shows an overview of the methodology. We created a mouse agent in the Unity game engine. The agent consisted of two cameras that were separated and oriented similarly to mouse eyes, and had a similar fields of view and resolution. We drove this agent through virtual environments to create mouse-perspective videos, and used these videos for self-supervised training of a deep artificial neural network model of mouse visual cortex. We created eight training datasets that were either naturalistic or non-naturalistic with respect to three factors: environment, camera optics, and mouse movement patterns. We used Representational Similarity Analysis (Kriegeskorte et al., 2008) to compare the stimulus-driven activations of model neurons with the responses of neurons recorded from the mouse visual cortex using two-photon calcium imaging (de Vries et al., 2020).

**Figure 1:**
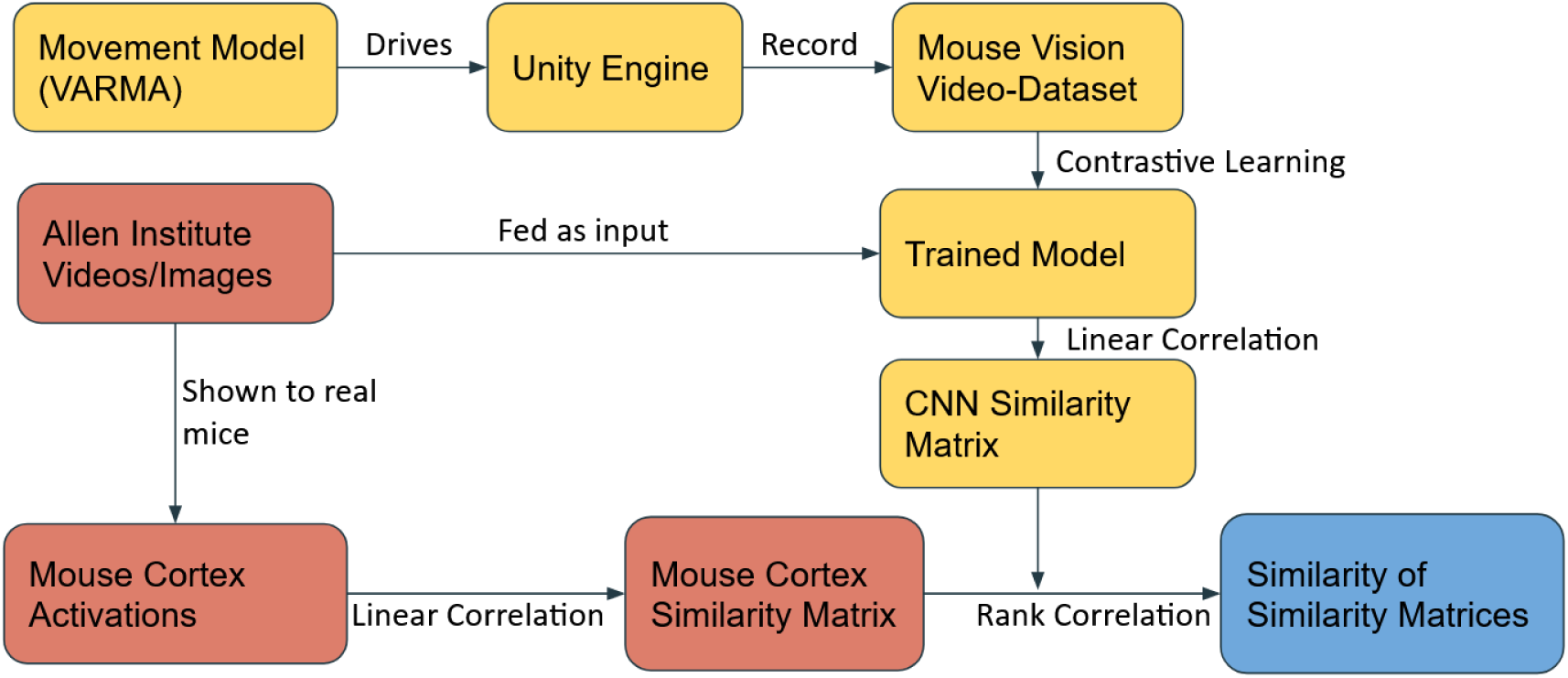
High-level outline of the methodology. The naturalistic model of motion was a vector auto-regressive moving average (VARMA) model based on mouse motion tracking data (Section 2.3)

### 2.1 Visual cortex model

We used MouseNet (Shi et al., 2022), a convolutional neural network that is based on mouse visual cortex anatomy. MouseNet is a feedforward two-dimensional convolutional model. It has layers corresponding to the dorsal lateral geniculate nucleus and layers 2/3, 4, and 5 of several areas of mouse visual cortex (VISp, VISl, VISrl, VISal, VISli, VISpl, VISpor). The model’s population sizes, kernel sizes, and spatial connection density profiles are based on neuroanatomical data. Following (Shi et al., 2022), the network’s output was produced by average-pooling and concatenating the outputs of cortical layer 5 of each cortical area in the model.

### 2.2 Training datasets

We created a mouse agent in Unity. The agent had two cameras with horizontal viewing angles of 135 degrees, separated by 1cm and 100 degrees horizontal visual angle, with the overlapping views forming a binocular region (Prusky et al., 2006). However, we only trained a monocular model corresponding to the right eye and left cortex. We recorded videos from this agent with a resolution of 64×64 pixels, approximating mouse visual resolution (Shi et al., 2022), and a frame rate of 50Hz. We recorded a total video length of 2 hours per dataset condition.

We created naturalistic and non-naturalistic versions of three simulation factors that affected the visual stimuli, specifically the virtual environment, the optical properties of the eyes, and the motion of the vitual mouse’s head through the environment. We experimented with all possible combinations, resulting in eight datasets (Table 1).

**Table 1:**
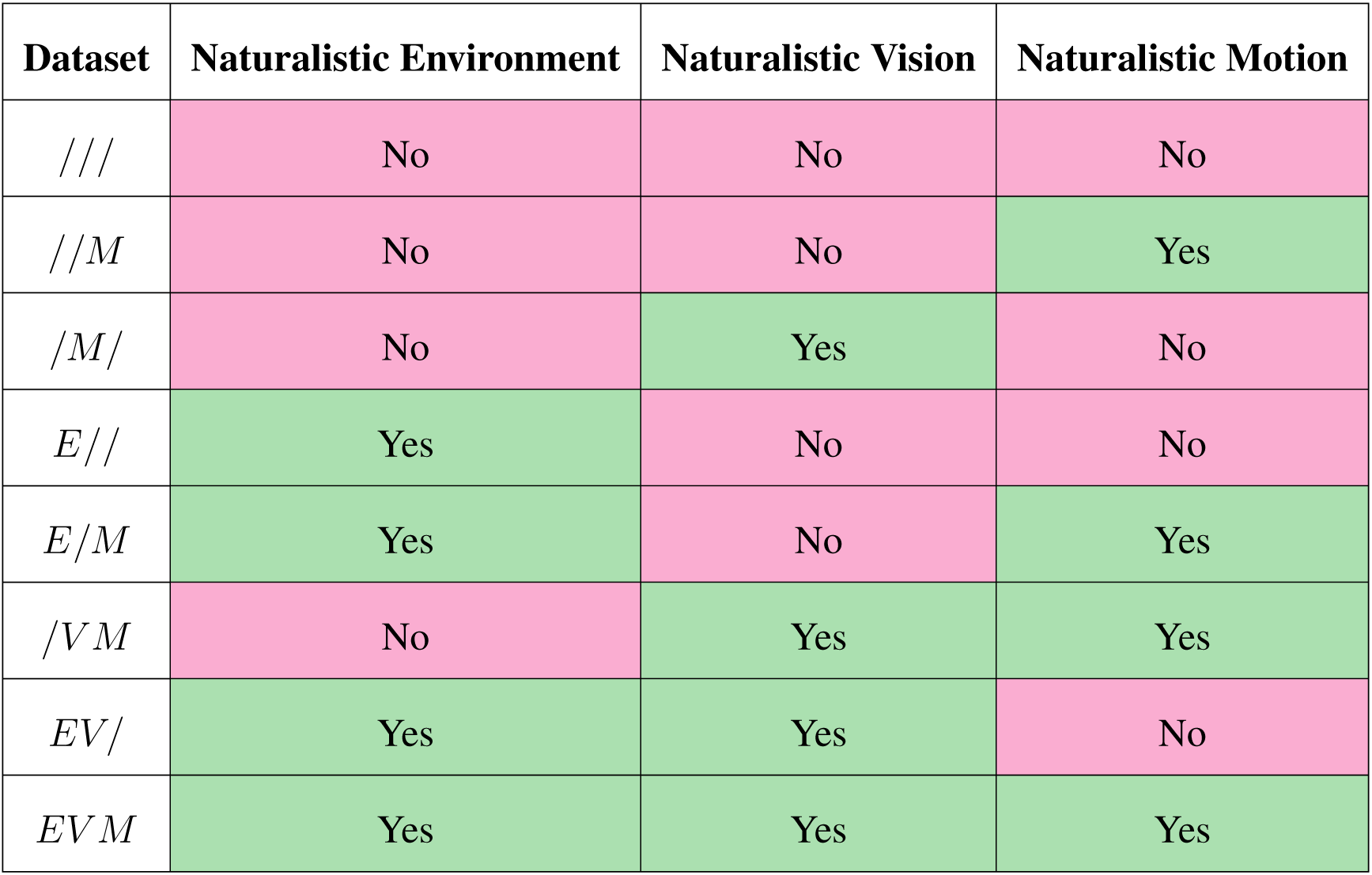
Overview of the datasets developed for this study. We developed eight video datasets that spanned all possible combinations of naturalistic and artificial models of three factors: the environment, eye optics, and pattern of motion of the mouse through the environment. Each dataset is identified with a three-character code indicating which factors were naturalistic. The letters E, V, and M indicate naturalistic model of the environment, vision, and motion, respectively. The character / in place of a letter indicates that a non-naturalistic model was used, as described in the text.

### 2.1 Environment

The realistic environment included swaying grass, trees, shrubs, a dirt path, rocks, apples, and logs (Figure 2). The artificial environment was modelled after a fictional spaceship, with grey metallic walls, floor, and ceiling. It had bright neon light strips on the walls and ceiling.

**Figure 2:**
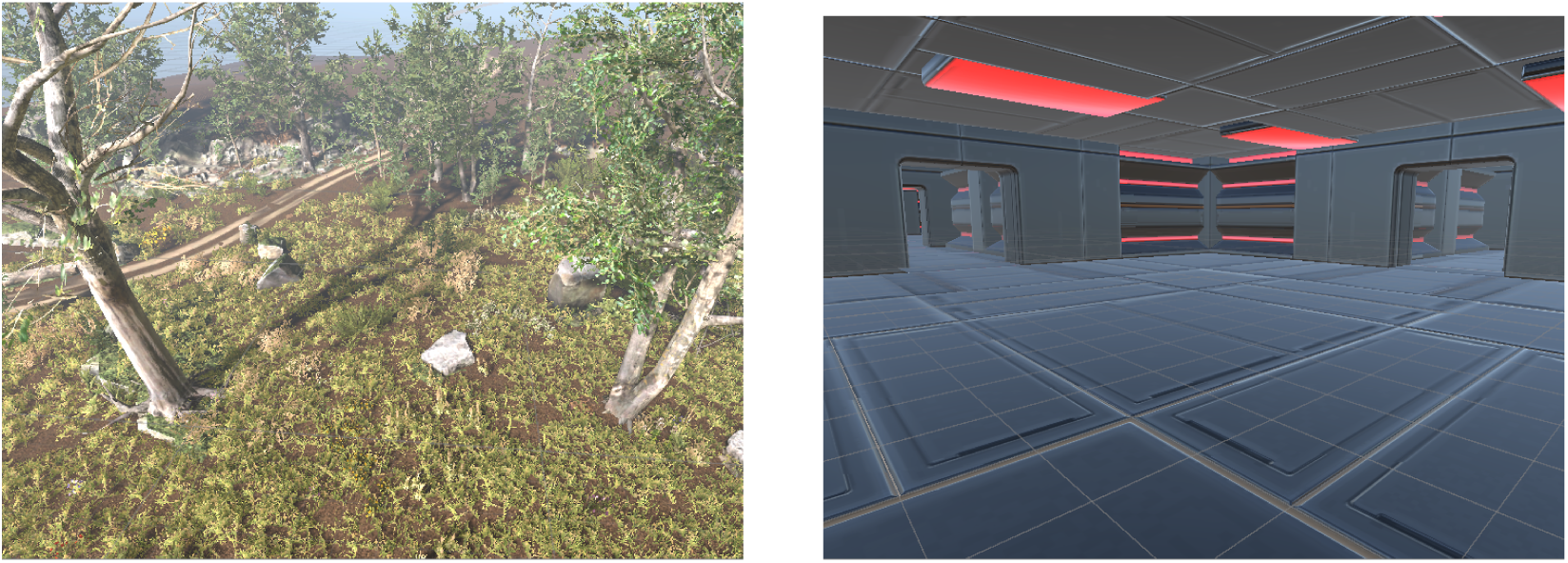
The naturalistic meadow environment (left) and the non-naturalistic spaceship environment (right). These broad views are not from the mouse model’s perspective.

### 2.2 Optics

We used a camera with a focal length of 2.6mm and numerical aperture of 0.49 as a naturalistic model of the optics of the mouse eye (Geng et al., 2011). Objects 10-15cm away were in focus, consistent with the near-sightedness of mice. Additionally, following (Shi et al., 2022), we averaged the red and green color channels to approximate mouse dichromatic vision. Artificial optics consisted of Unity’s default pinhole camera, accompanied by red, green, and blue color channels.

**Figure 3:**
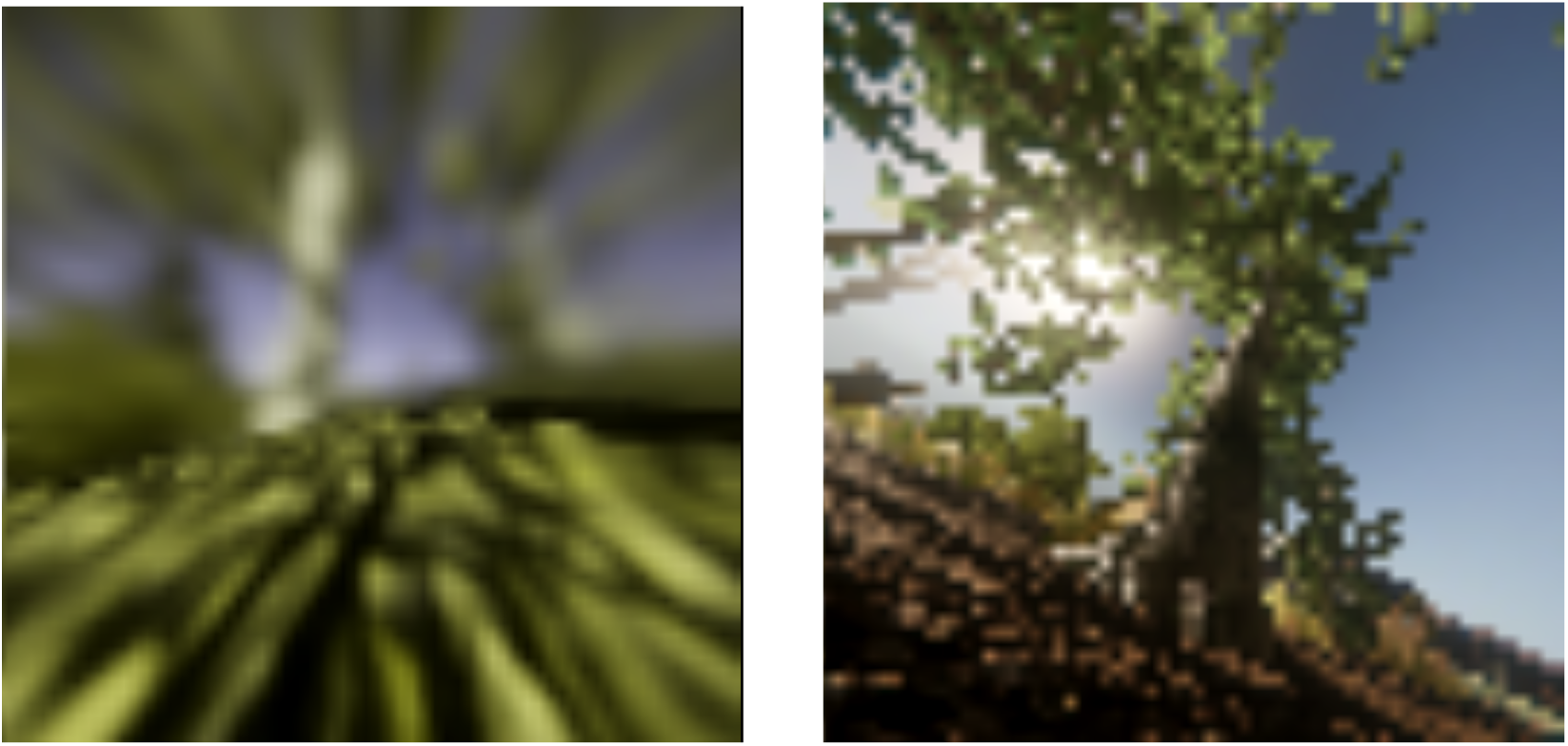
Images collected from Unity with the naturalistic (left) and artificial (right) optics models.

### 2.3 Mouse motion

The artificial motion model consisted mainly of forward motion in a straight line, punctuated by left and right turns. Straight-ahead motion continued for 100-200 frames at a time, at 50Hz, and turns were 45-135 degrees. Both forward duration and turn magnitude were uniform random variables. Turns occurred over 10 frames.

The naturalistic model was derived from mouse motion tracking data. We used data from the Peyrache lab, from a mouse that moved freely in a 50cm x 50cm box for 30 minutes while its head position and orientation were recorded at 100Hz. We developed a statistical model of this dataset, allowing us to generate naturalistic trajectories of arbitrary duration through the larger Unity environments. Specifically, we developed a Vector Auto-Regressive Moving Average (VARMA) model, which acts like an infinite impulse response filter with white noise as input.

Qualitative analysis of the data revealed two distinct motion patterns in which the mouse was ambulating or not. We hand-labelled these two states throughout the dataset and modelled them separately, focusing on longer state durations of 8s or more and ignoring brief state fluctuations. We then fit shifted exponential distributions (beginning at 8s) separately to the durations of each state, and sampled from these distributions to switch between models at generation time. The scale parameters were 14.5s and 18.1s for ambulating and non-ambulating, respectively. To switch smoothly between models, we introduced a seven-frame blank interval and interpolated.

**Figure 4:**
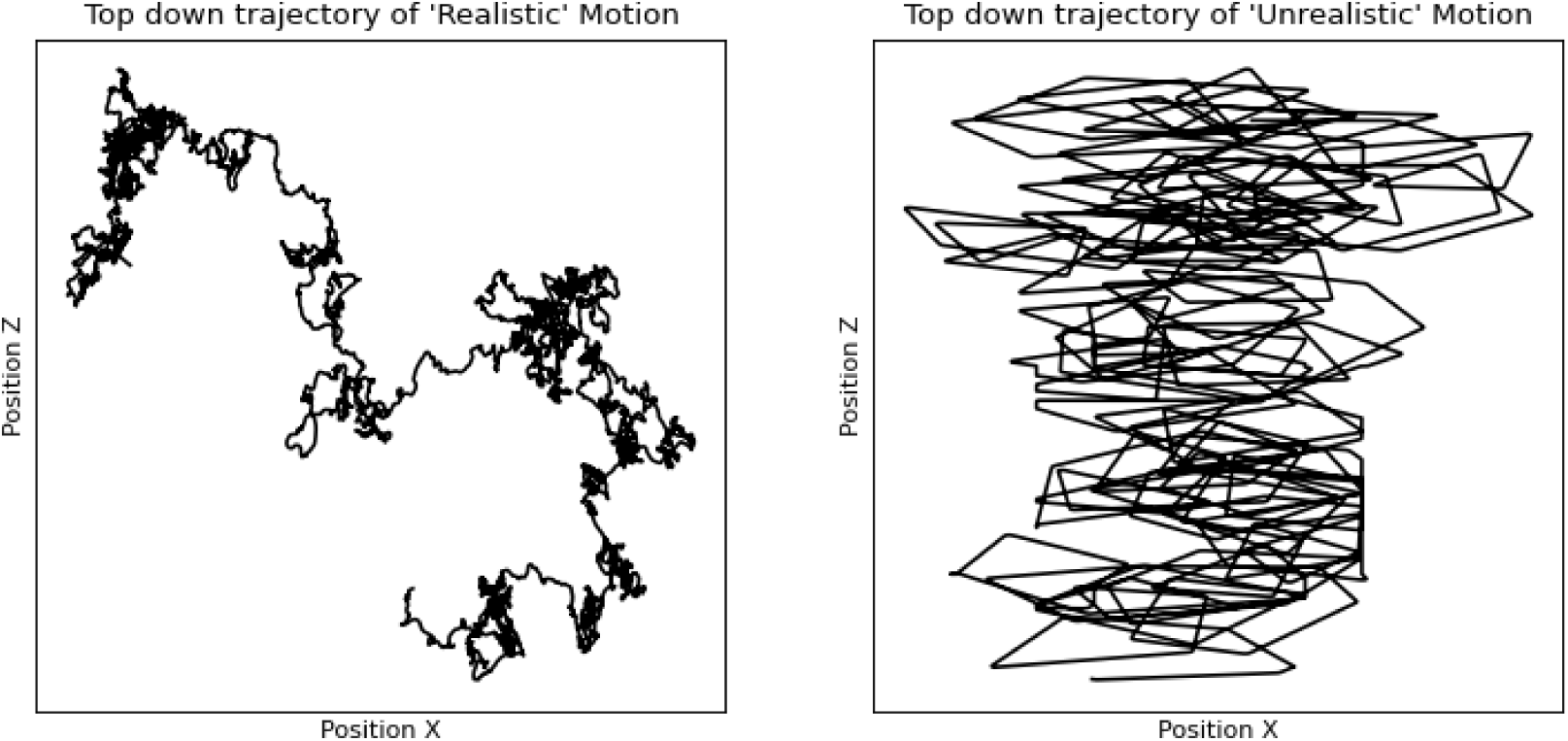
Comparison of realistic motion (left) with non-realistic motion (right) from a bird’s eyed view.

The motion-tracking data included three-dimensional global position coordinates and three Euler angles of the mouse head. Before modelling the data we processed it to reduce noise. This processing was done with the angles in quaternion space to avoid discontinuities. Quaternions represent each orientation redundantly on opposite sides of a four-dimensional unit hypersphere, so that *Q* and −*Q* represent the same physical angle. We flipped quaternions as necessary so that consecutive quaternions were never separated by an angle *> π/*2 on the hypersphere. This resulted in a smooth and continuous representations of orientation.

To reduce the impact of sensor noise, we discarded samples in which the quaternion angle changed by more than 0.125 radians in one 10ms step. We chose this maximum by inspecting a histogram of rotation speeds in the head-motion data. Discarded orientation samples were replaced by spherical linear interpolation (Shoemake, 1985).

Each channel of motion data was then low-pass filtered backwards and forwards with a 25Hz 5th-order Butterworth filter, then down-sampled by a factor of two in order to match the frame rate of the Unity models. The quaternions were then normalised back onto the unit hypersphere, as linear filtering introduced slight deviations.

We modelled the mouse’s horizontal velocity, rather than position, to allow generation of trajectories over a larger area. However, we modelled the vertical position directly, so that the elevation above the ground would remain in a realistic range. We also converted absolute velocities to egocentric velocities (based on head angle). Similarly, we modelled pitch and roll angles directly because they are bounded during standing and locomotion, but we modelled yaw velocity to allow continuous changes in heading direction.

We used a vector autoregressive moving average (VARMA) model to produce random trajectories that resembled mouse head motion. Input to the VARMA model consisted of a 6D white noise signal, *ɛ*. The hyperparameters of a VARMA model are *p* and *q*, which correspond to the autoregressive and moving-average orders, respectively, i.e. the number of past output and input samples that affect the current output. The learned parameters are the coefficients *ϕ_i_* and *θ_i_* in,

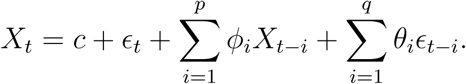

The VARMA models were trained using the statsmodels Python package Seabold and Perktold (2010). To choose the hyperparameters, we performed a grid search over *P, Q* ∈ [1*…*9] (chosen by inspection of preliminary experiments). We chose the parameter combination that optimized the Akaike Information Criterion. This resulted in (*p, q*) = (7, 2) for the ambulating state and (*p, q*) = (5, 5) for the non-ambulating state.

We sanity-checked the resulting models by manually comparing cross-correlations between dimensions of recorded and generated trajectories (Figure 17 in the Appendix), as well as univariate distributions of each trajectory dimension, and pairwise bivariate distributions (Figure 18), and finally by watching an animated mouse in Unity which followed generated trajectories.

### 2.3 Synthetic Environments

We suspected that differences in brain similarity due to training in the meadow and spaceship environments were due to the simple statistical properties of the corresponding videos. To test this hypothesis, we created new Unity environments and training videos that approximated the E// and /// conditions in terms of spatial frequency content, edge-orientation distribution, colour distribution, and temporal autocorrelation.

The new environments consisted entirely of cubes and spheres of various colours and sizes (Figure 5). The synthetic spaceship scene also included a visible ground plane. The synthetic meadow did not, because the meadow ground colour distribution was diverse. The shape sizes were drawn from six bins with size scaling uniformly sampled from the ranges 1-5, 5-10, 10-25, 25-50, 50-75, and 75-100. We optimized the number of objects drawn from each bin and the ratio of spheres to cubes to approximate the 2D spatial frequency spectra of the meadow and spaceship environments. These spectra encapsulate the environments’ mean spatial frequency rolloffs and edge-orientation distributions. The final approximations of these spectra are shown in Figure 6.

**Figure 5:**
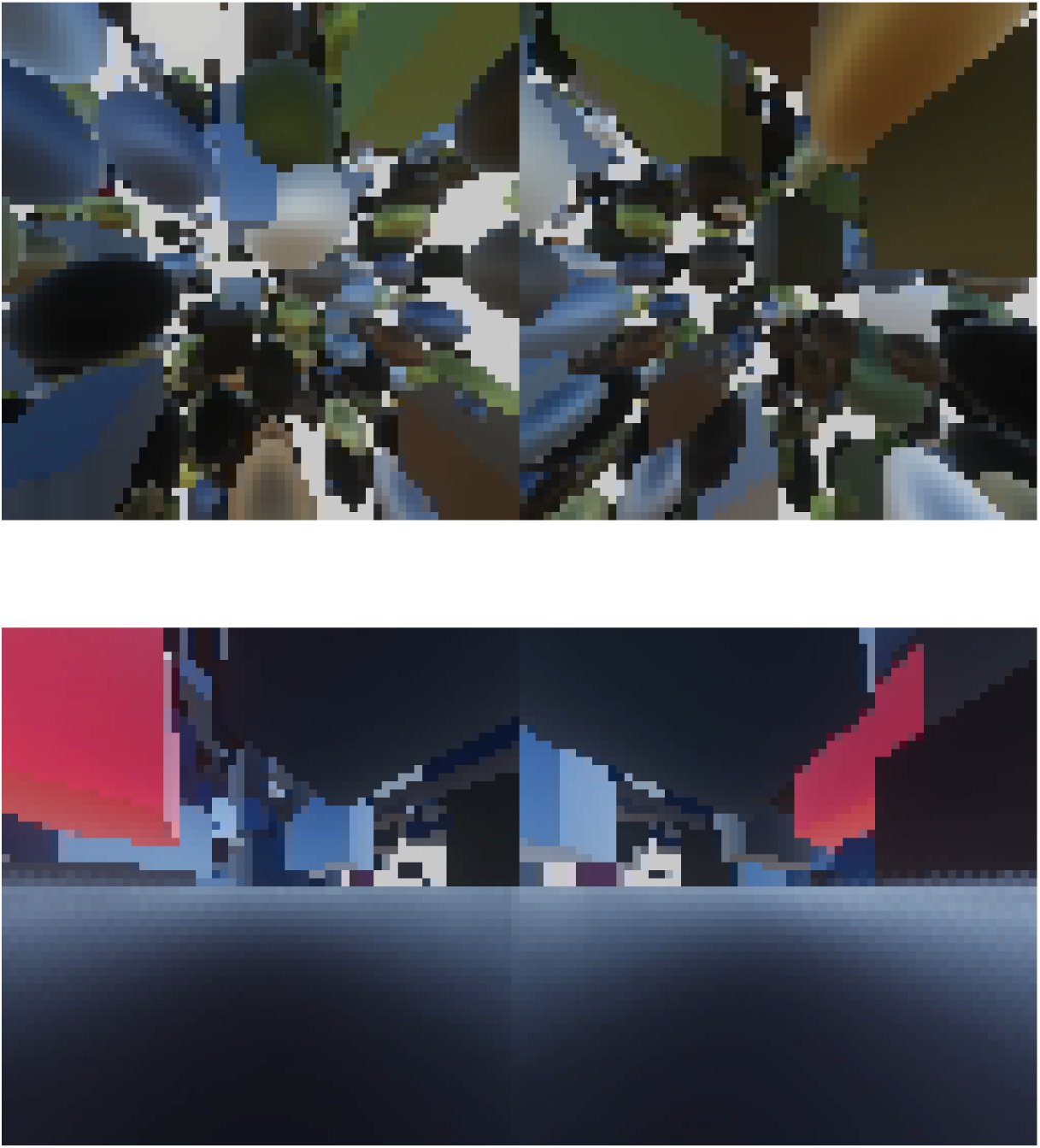
Example scenes from the synthetic meadow (upper) and spaceship (lower) environments as can be seen from the left and right mouse eye.

**Figure 6:**
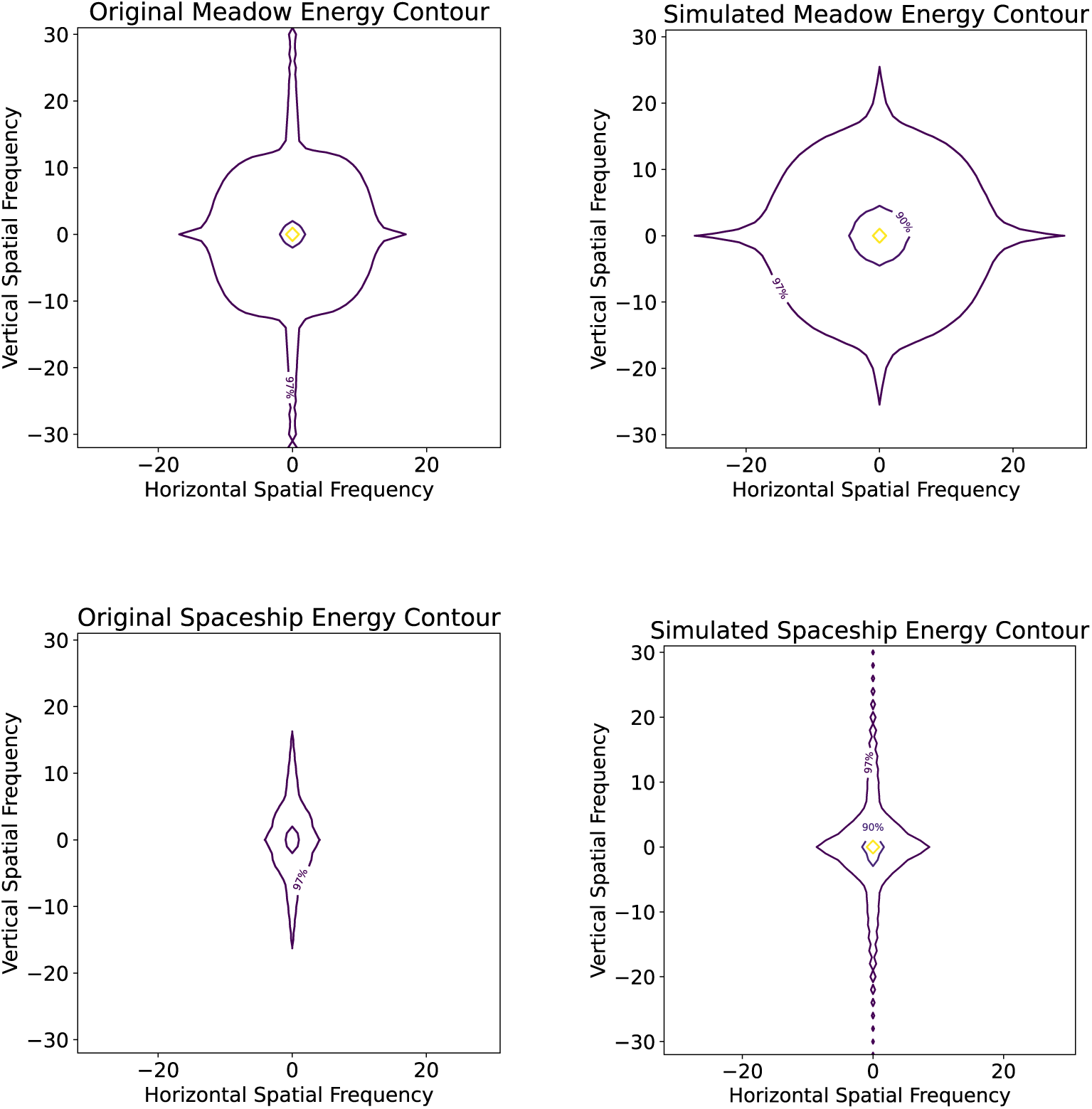
Spectral energy contours at the 80%, 90%, and 97% thresholds.

Each environment can have an object density between 0 to 600 objects per bin within a 1000^3^ cubic unit volume with up to a maximum of 3600 objects within the boundary of the scene. The parameters were optimized via random search using 10,000 videos to find closest matches to the spectra of the original spaceship and meadow scenes. Each of the recordings contained 50 random frames sampled using the same parameters. The weighted mean and standard deviation of the top five parameter sets resulting in the lowest mean squared error between the spectrum decay rates, averaged over 63 orientations interspersed between the vertical and horizontal directions were calculated and can be seen in Table 2. The table shows significant bias towards high ratio of spheres for the meadow environment and cubes for the spaceship environment.

**Table 2:** Weighted mean and standard deviation of the top five parameter sets which results in the closest matching spaceship and meadow spectra signature.

| Bin Sizes | 1-5 | 5-10 | 10-25 | 25-50 | 50-75 | 75-100 | Percent of Spheres |
| --- | --- | --- | --- | --- | --- | --- | --- |
| Meadow |  |  |  |  |  |  |  |
| mean | 253 | 239 | 404 | 276 | 393 | 445 | 87% |
| std | 148 | 194 | 145 | 205 | 198 | 208 | 8% |
| Spaceship |  |  |  |  |  |  |  |
| mean | 275 | 162 | 341 | 416 | 502 | 566 | 3% |
| std | 194 | 130 | 188 | 195 | 66 | 17 | 2% |

We set the colour of each sphere/cube to match a randomly drawn pixel from the meadow or spaceship video. We inspected histograms of the resulting color-channel (red, green, blue) brightness values and found that they were not as smooth as those of the meadow and spaceship environments. To improve smoothness, we used Unity’s default shader to introduce lighting-related variations across the surface of each shape. As a final step, we matched the brightness histograms of each colour channel by post-processing the recorded videos. We did this by mapping the synthetic histogram distributions to the meadow/spaceship distributions as *b^′^* = *F^−^*1(*F_s_*(*b*)), where *b* is the synthetic channel brightness, *F_s_* is the cumulative density function of the synthetic-video distribution, and *F_o_* is the cumulative density function of the original (meadow/spaceship) distribution.

Finally, we modulated the speed of the mouse through the new environments to match the average pixel-wise autocorrelations of the meadow/spaceship videos. We minimized the mean difference between the autocorrelation functions of the original and synthetic videos over lags of 1-40 frames (corresponding to lengths of video segments used in model training). We used the method of false position to find the closest fit numerically. The best approximations are shown in Figure 7 with 77x for the simulated meadow environment and 67x for the spaceship.

**Figure 7:**
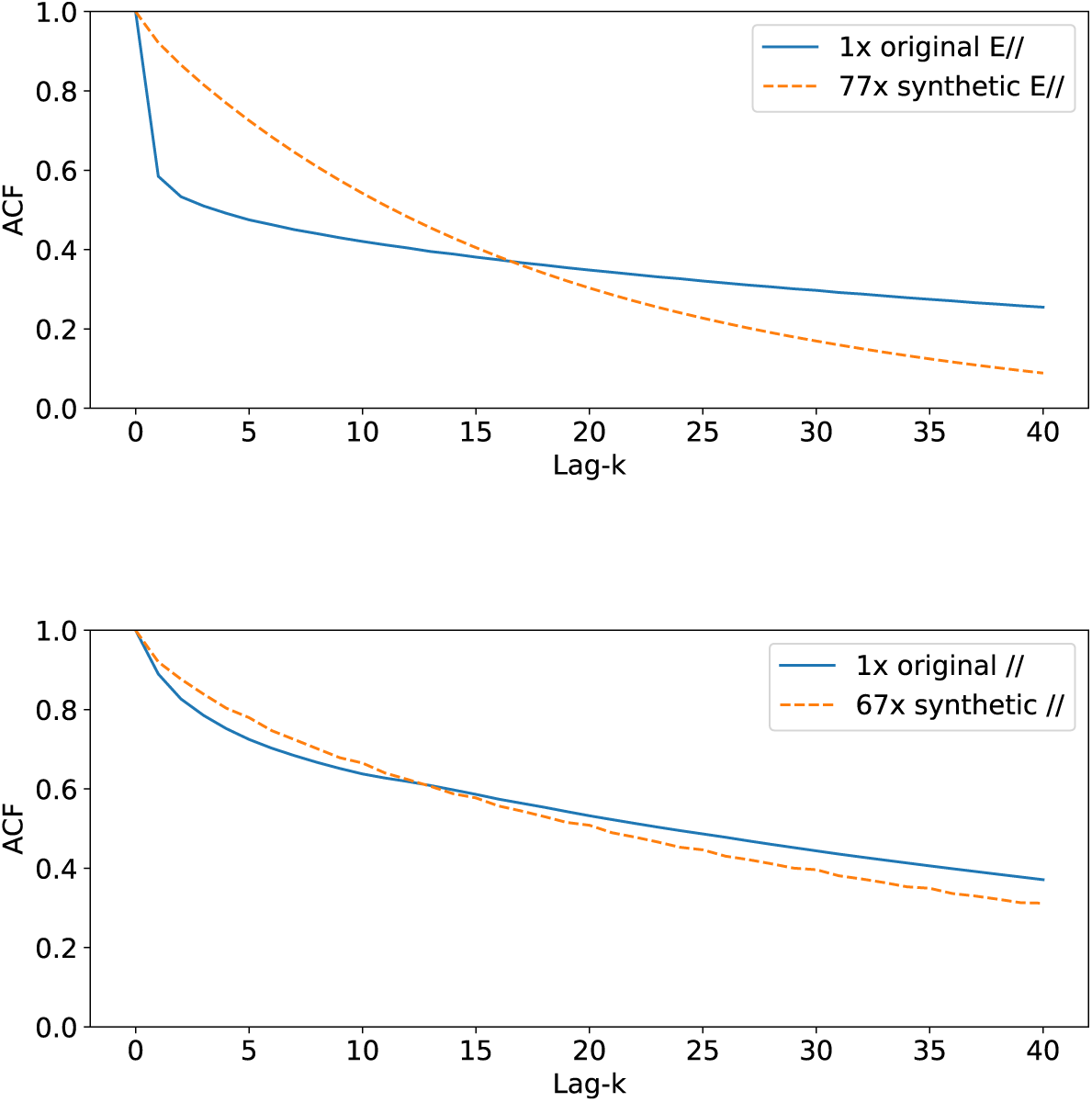
Autocorrelation of individual pixel values over the lags from 0 to 40 frames for meadow (top) and spaceship (bottom) scenes.

### 2.4 Training procedure

We trained MouseNet on each video dataset using the self-supervised Dense Predictive Coding (DPC) (Han et al., 2019). The goal of DPC is to accurately identify latent representations of future video frames from a random batch of distractors, using the current and past frames. Following (Han et al., 2019) we added a recurrent head to MouseNet, consisting of a convolutional gated recurrent unit. This head received inputs from layer-5 of each cortical area in MouseNet, and downsampled by average pooling to 4×4 resolution. It had 144 input channels, one hidden layer with 144 channels, input and recurrent kernel sizes of 1×1, and dropout with 10% of channels dropped at random. It compressed multiple frames into context variable that was used to predict embeddings of future frames.

We divided training videos into sequences of eight five-frame clips. Sequences were organized into random mini-batches of size 32. The batch size was limited by 32GB memory on Nvidia V100 graphical processing units. The model used the first 5-7 clips (25-35 frames) in each sequence as context, and was trained to select the subsequent five-frame clip from among negative examples in the same mini-batch.

For each training dataset condition, we trained the model five times with different random initializations.

### 2.5 Comparison with brain data

We compared the outputs of model neurons with public two-photon calcium imaging data from the Allen Brain Observatory (de Vries et al., 2023). In particular, we used data recorded from mouse visual cortex while the mice watched 30s natural movie clips. We selected the CEX2-CRE2 Cre line, which was the line with the highest noise ceiling. Data from this Cre line involve areas VISp, VISl, VISal, VISpm, VISam, and VISrl. However, we omitted VISrl from the analysis because it had a very low noise ceiling. These recordings come from cortical depths between 175-250 µm, and 265-250 µm, corresponding roughly to cortical layers 2/3 and 4, respectively. We used spike-rate estimates from events in non-overlapping 30ms windows.

We quantified alignment between the models and mouse visual cortex using Representational Similarity Analysis. First, we input the same natural movie clips to the MouseNet models as were shown to the mice. We then calculated similarity matrices for each layer of the model and each mouse cell population. These matrices consisted of Pearson correlations between the responses to pairs of 180 five-frame stimulus clips. Brain similarity scores were then calculated as Kendall’s *τ* rank correlations between pairs of model and mouse similarity matrices (Nili et al., 2014).

We normalized similarity scores relative to inter-animal noise ceilings. We calculated noise ceilings as mean similarity scores for each cell population between data from 100 randomly selected pairs of mice. The noise ceilings ranged from 0.30 to 0.63. We report similarity scores as a fraction of the corresponding noise ceilings.

Due to the lag between stimulus presentation and neural response in mice, we compared model responses to mouse brain responses two frames (67ms) later. Responses across the mouse visual cortex are strong at this lag (Siegle et al., 2021). We found that the results were not highly sensitive to lags of 0-10 frames, perhaps due to autocorrelations in the video stimuli.

## 3 Results

### 3.1 Impact of stimulus properties on latent prediction accuracy

The models across all training conditions developed acceptable prediction accuracies that were well above chance. Top-1 training accuracies with different datasets ranged from roughly 0.4 to 0.7 after 100 epochs. This wide range of accuracies is expected because some datasets required more difficult predictions. Unsurprisingly, predictions were most accurate (∼0.65-0.7) in the datasets that combined artificial environment and artificial motion. Distinguishing next frames in these datasets was probably more straightforward, because optic flow followed a consistent expansion pattern, and certain visual features were distinctive. Predictions were least accurate (∼0.4) in conditions that combined naturalistic environment and motion.

### 3.2 Impact of stimulus properties on mouse brain alignment

Figure 8 shows brain similarity scores for each training data condition, averaged over brain areas and random seeds. The EVM condition (all factors naturalistic) led to the highest brain similarity scores. Conditions with the naturalistic environment produced much higher similarity scores than those without. The /V/ condition produced slightly lower similarity scores than the /// condition. Otherwise, conditions with any combination of naturalistic factors resulted in higher scores than ///.

**Figure 8:**
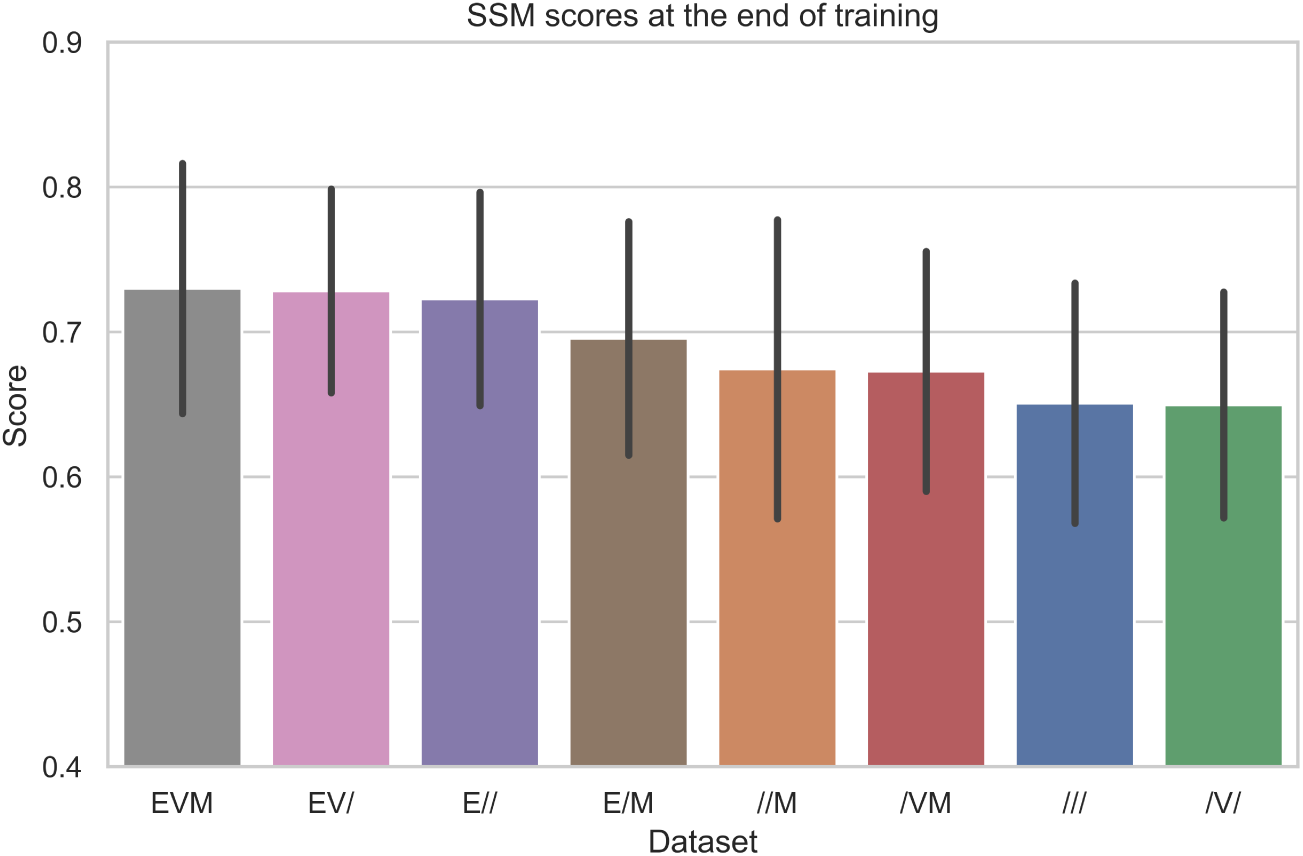
Maximum similarity scores across CNN-areas for each mouse area at the end of training, averaged across mouse cell populations and random seeds. Error bars show standard deviation across both mouse-areas and random seeds.

Figure 9 summarizes similarity with mouse brain activity across mouse cell populations and training conditions. Trained models generally had greater similarity with mouse data than untrained models, across cell populations, except //M models relative to mouse VISal 2/3 and /v/ models relative to VISam 2/3. Similarity with most mouse cell populations was higher in conditions with the natutalistic envionment, but this was not true for VISp 2/3.

**Figure 9:**
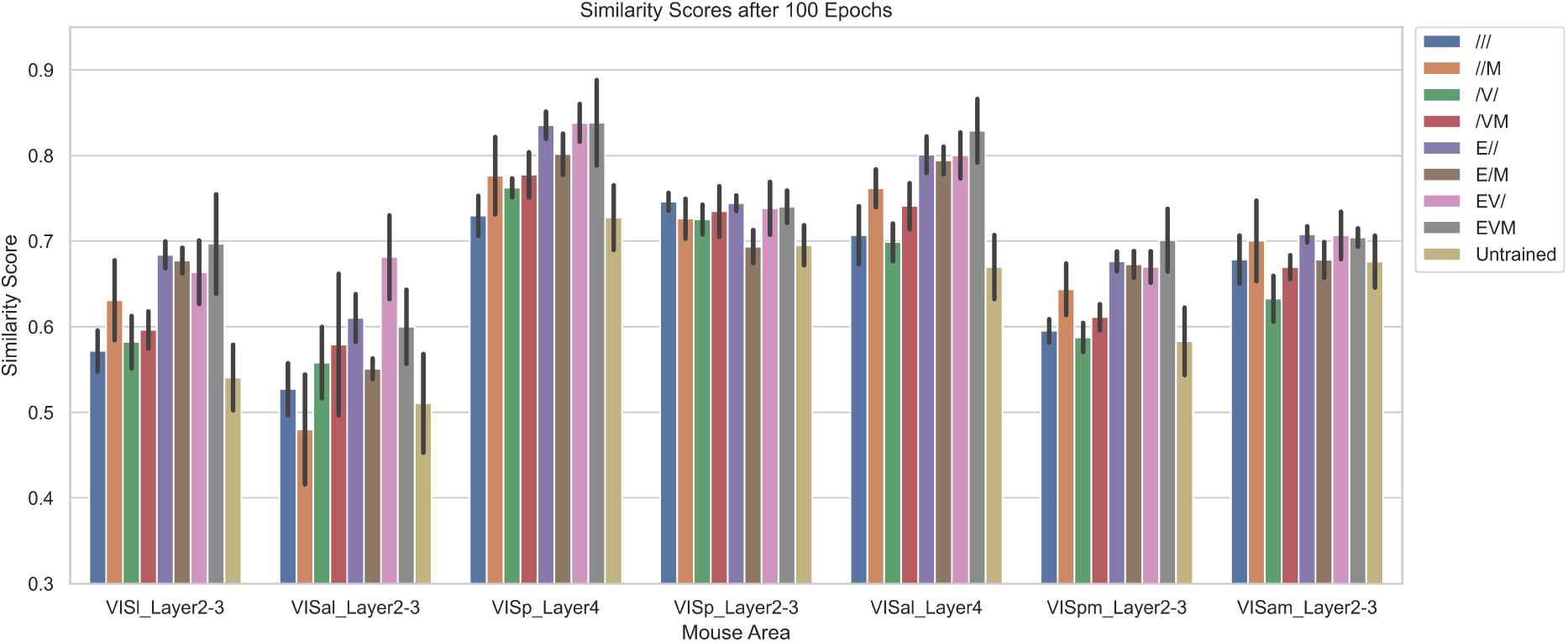
Similarity of model activity with different mouse cell populations (maximum for each mouse area across model areas), in different training conditions, after 100 epochs of training. The error bars show mean +/-standard deviation across five random initializations.

An ANOVA revealed a significant main effect of the environment condition (*p <* .0001). There were also significant two-way interaction effects between environment and optics (*p* = .0001), environment and motion (*p* < .0001), and optics and motion (*p* = .0017), and a significant three-way interaction effect (environment, optics, and motion; *p < .*0001). Therefore, differences between these training stimuli significantly impacted brain alignment, and the impact of any given factor depended on other factors.

We performed regression analysis to quantify the average impact of various training conditions relative to the baseline with all unrealistic properties. We used binary vari-ables reflecting the seven cell populations and seven remaining training conditions as regressors (Table 3). Conditions with the naturalistic environment resulted in substantially greater brain similarity than those with the artificial environment. In particular, the regression coefficients for the E/M, E//, EV/, and EVM conditions were 0.045, 0.072, 0.078, and 0.0792, respectively (all significant with *p < .*001). Brain similarity after training with the /// dataset averaged 0.650 across cell populations, so these are relative increases of 6.9% to 12.1%.

**Table 3:**
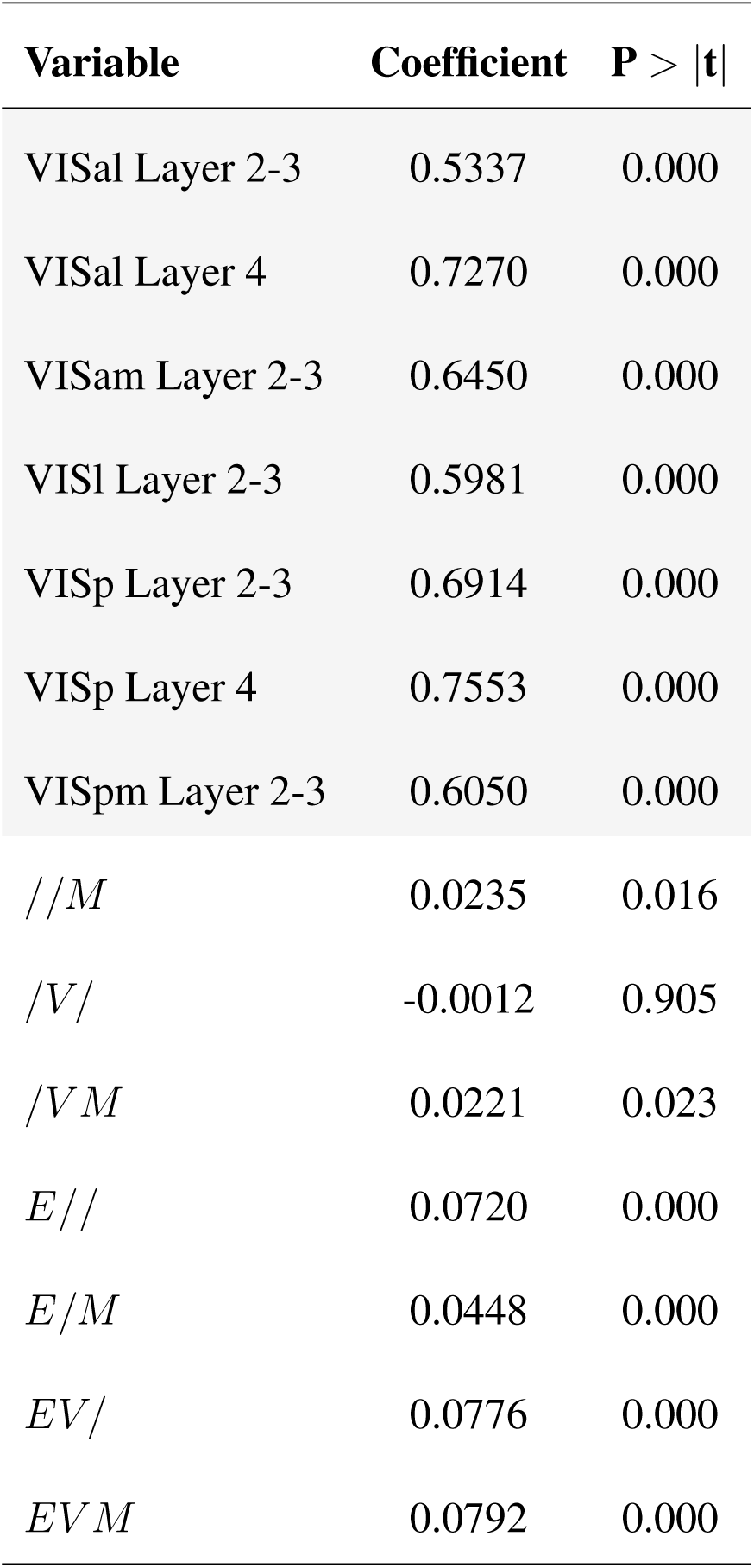
Regression analysis of various dataset training conditions. Coefficients for the dataset conditions (unshaded) represent change in similarity with a given mouse area (shaded) compared to baseline /// condition.

The naturalistic optics models had minor positive and negative impacts in different conditions, with an almost neutral impact on average. The greatest impact was in conjunction with naturalistic environment and motion, leading to brain similarity 0.0344 greater (from the regression model) in the EVM condition vs. the E/M condition. However, on average across conditions, naturalistic optics had little impact on brain similarity (+0.009). Across conditions, naturalistic optics consistently increased similarity with VISal Layer 2/3 and VISp Layer 4, with inconsistent effects otherwise (Figure 9).

Naturalistic motion also had a small average impact on brain similarity (+0.0053 on average, from the regression model), with inconsistent effects across mouse areas and conditions. VISl Layer 2/3, VISal Layer 4, and VISpm Layer 2-3 all showed the same pattern of increased similarity owing to naturalistic motion (Figure 9). In these areas, the average increase owing to naturalistic motion was 0.02666, which is a 4% relative increase. In other mouse areas the impacts of naturalistic motion were also substantial, but in different directions across pairs of conditions (e.g. /// vs. //M and /V/ vs. /VM).

### 3.3 Time course of changes during training

Self-supervised prediction loss decreased and accuracy increased with monotonic trends during training, without saturating after 100 epochs (Figures 10 and 11). Among conditions with the realistic environment, those with realistic motion led to a more rapid early drop in loss and corresponding rise in accuracy. This is surprising, because realistic motion speed and direction varied on short timescales, which would be expected to make future frames harder to predict. However, accuracy was lower in these conditions late in training.

**Figure 10:**
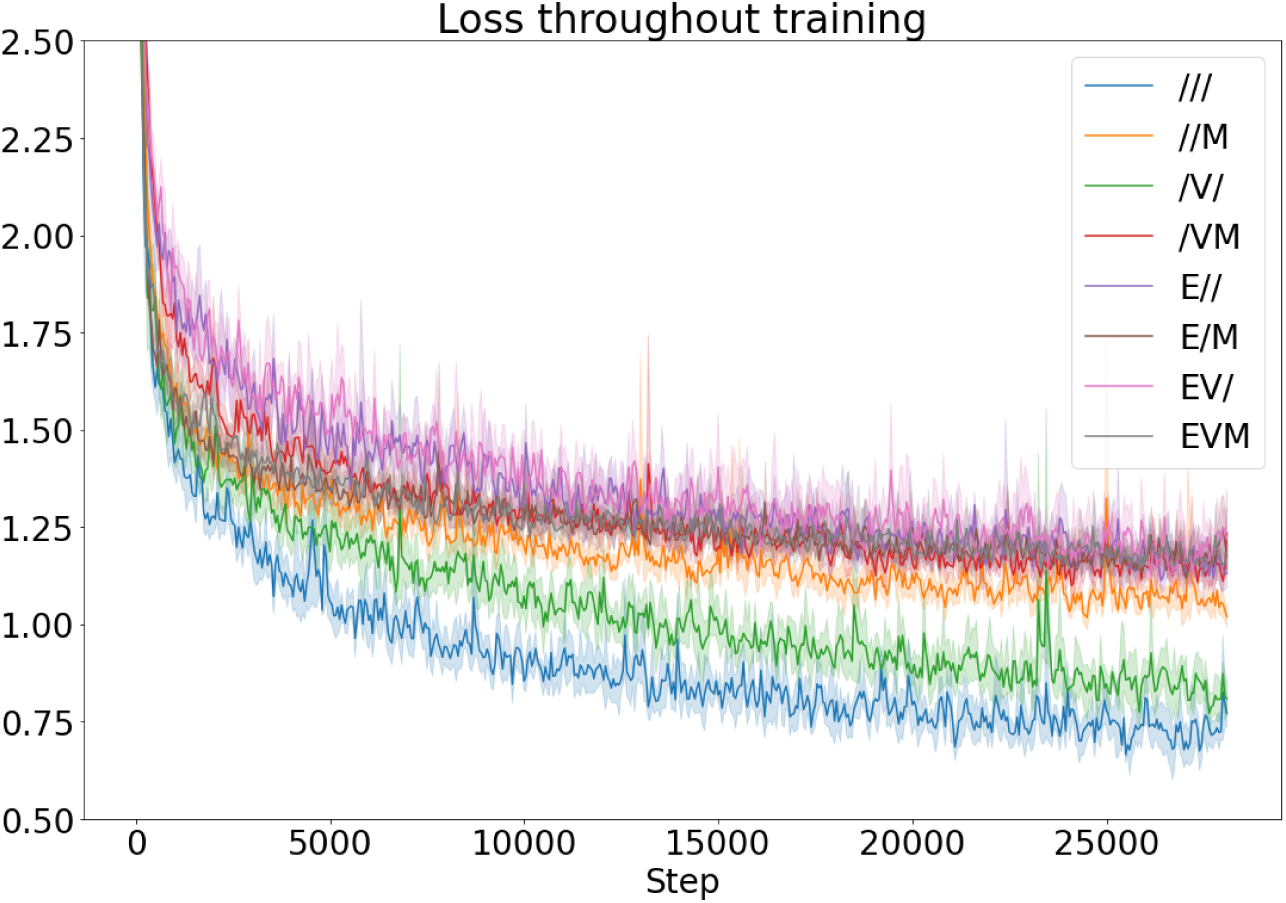
Training loss. Each trace shows mean (solid line) over five runs with dif-ferent random parameter initializations, smoothed with kernel [.6, .4]. The shaded area shows the 95% confidence interval over runs. The 28100 training steps (minibatches) correspond to 100 epochs.

**Figure 11:**
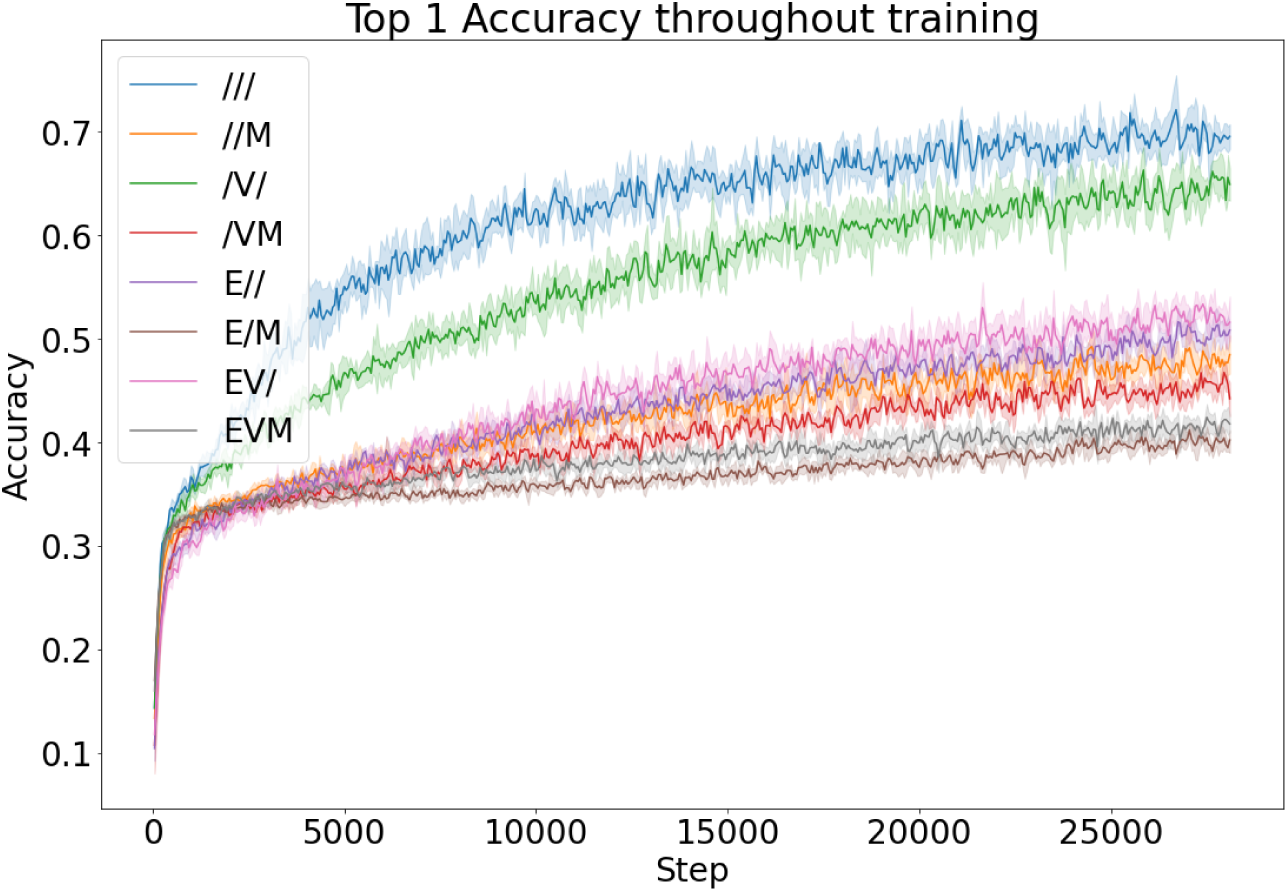
Top-1 training accuracy. Conventions as in Figure 10.

Mouse brain similarity did not develop monotonically, and its time course during training varied substantially across training conditions and cell populations. As an example, Figure 12 shows how similarity with VISal Layer 2/3 cells developed in the different training conditions. In most conditions there was a rapid drop in similarity at the beginning of training, followed by gradual and essentially monotonic increases or decreases in different conditions. Similarity had saturated after 100 epochs in most conditions, except perhaps in the EV/ and EVM conditions.

**Figure 12:**
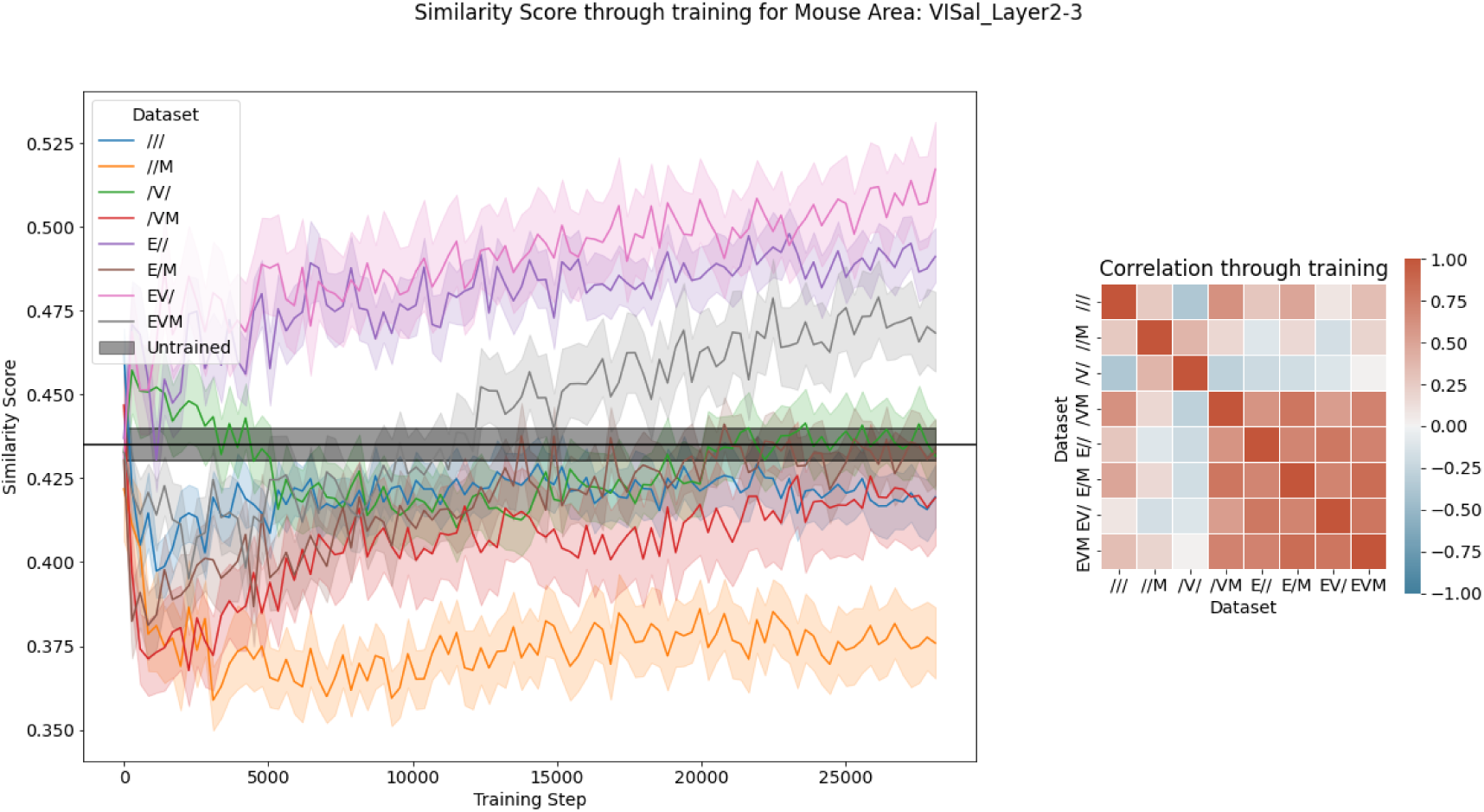
Left, Evolution of model similarity with VISal Layer 2/3 over the course of training (mean and 95% confidence interval for each condition over five models trained with different random initializations, and over model layers). The dark shaded area indicates the mean and 95% confidence interval of similarity over 28 untrained models with different random initializations. Right, Pairwise correlations of the training curves of different conditions.

As a contrasting example, Figure 13 shows similarity curves with respect to VISam Layer 2/3. Untrained models had relatively high similarity with these cells, and training reduced the mean similarity (across model layers and initializations), with the exception of //M. Changes in similarity with these cells developed primarily in the first 20 epochs.

**Figure 13:**
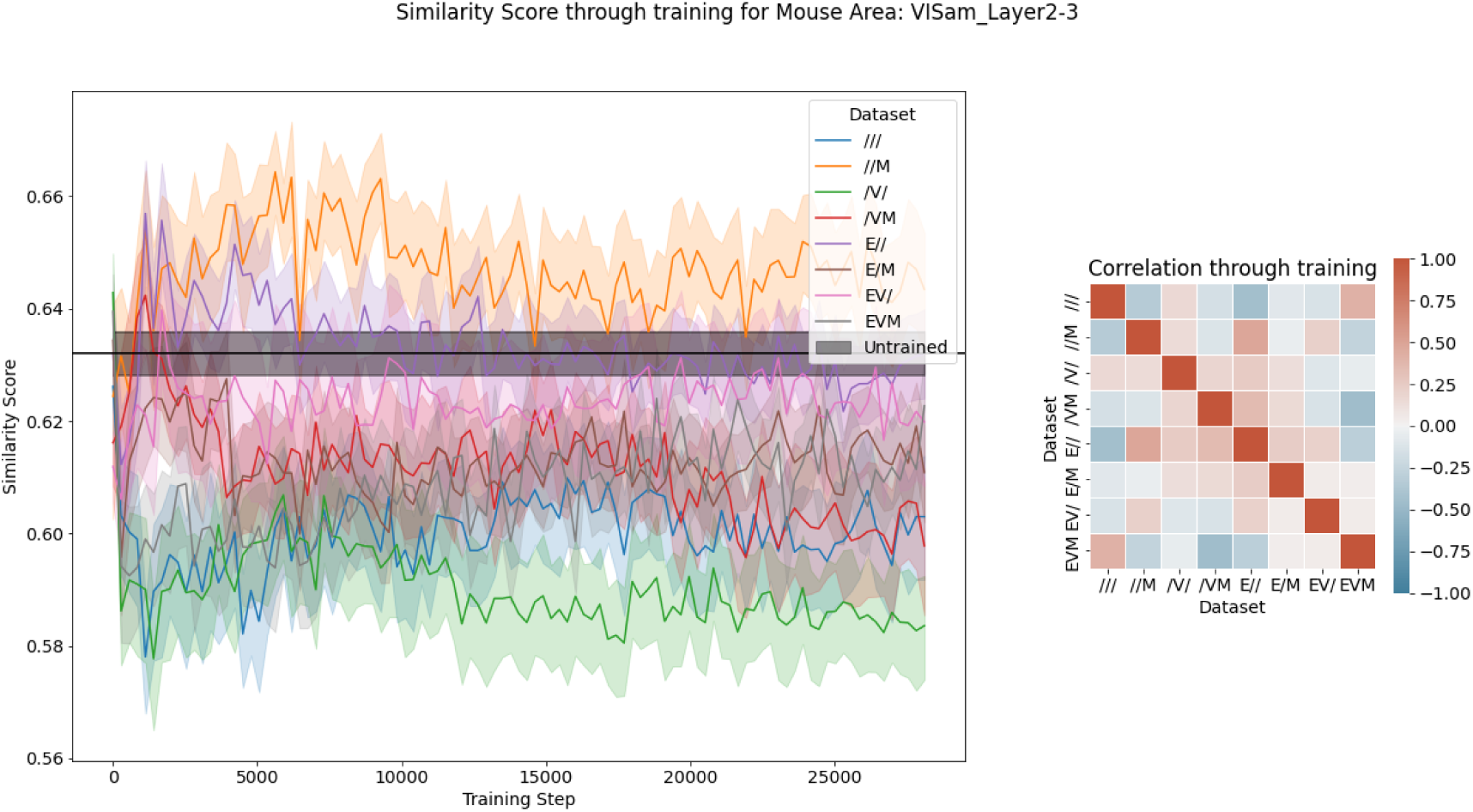
Evolution of mean model similarity (over initializations and model layers) with VISam Layer 2/3 over the course of training. Conventions as in Figure 12.

Across other cell groups (not shown), the vast majority of similarity curves changed quickly over the first few epochs, and saturated by the end of training, despite continued improvements in self-supervised prediction loss.

### 3.4 Relationship between self-supervised prediction loss and brain similarity

Across training conditions, there was a positive correlation between prediction loss and brain similarity, with high loss in the realistic-environment conditions that lead to high brain similarity. The *///* and */V/* conditions resulted in particularly low loss and high accuracy, and these models also had the lowest brain similarity overall. Conditions with more difficult predictions may have required more sophisticated representations, resulting in better brain alignment.

Within each training condition, across training steps and random seeds, self-supervised prediction loss and brain similarity were weakly related (Figure 14).

**Figure 14:**
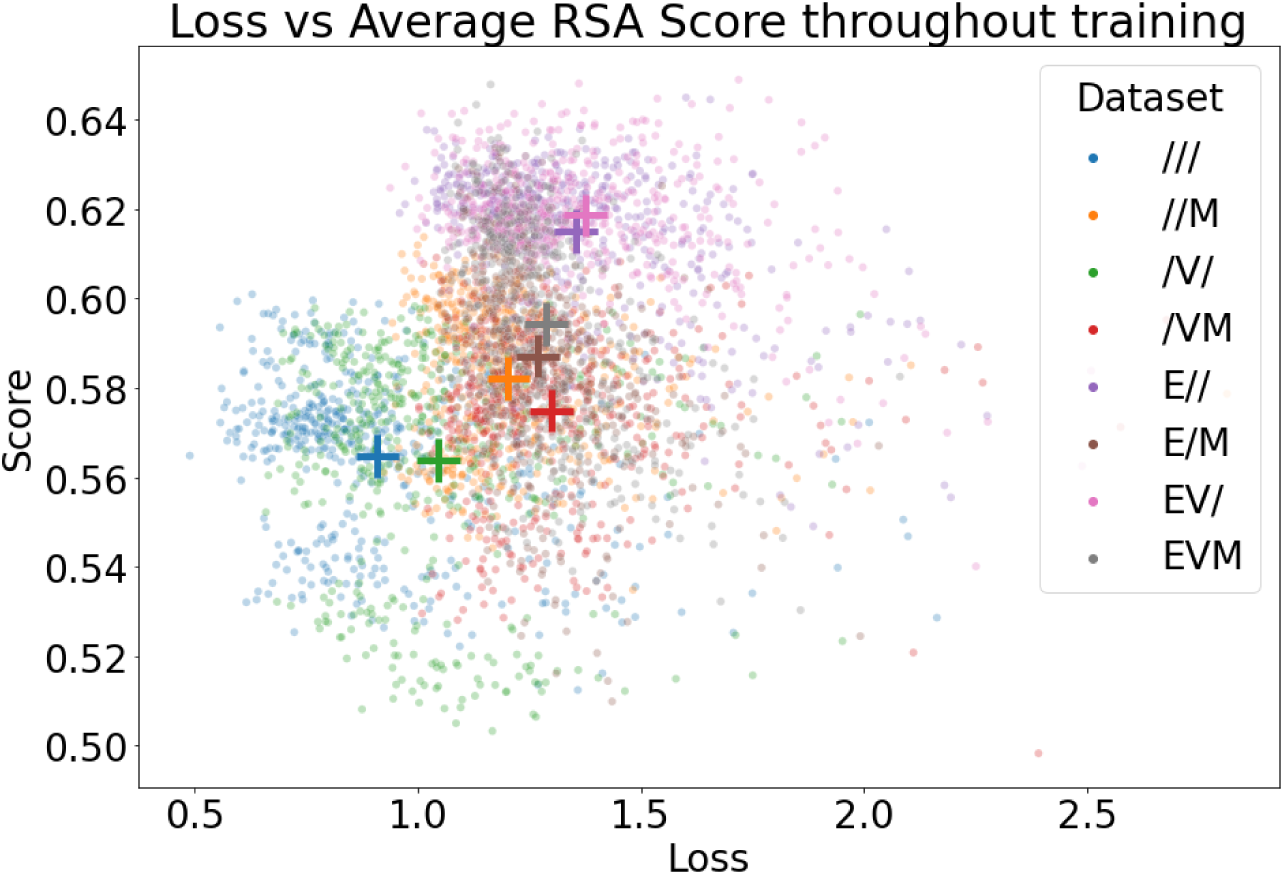
Representational similarity vs. training loss at each epoch after each random initialization. ‘+’ Markers represent the average similarity score and loss throughout training and across initializations.

### 3.5 Synthetic environments

To better understand which properties of the meadow vs. spaceship environment accounted for differences in brain similarity, we used abstract synthetic environments (made of cubes and spheres) that approximated some of their statistical properties. In particular, the synthetic environments approximated the original environments’ 2D spatial frequency spectra (which includes information about orientation distributions), colour distributions, and temporal autocorrelations (Section 2.3).

Unexpectedly, these properties did not account for brain similarity differences be-tween the meadow and spaceship environments. Figure 15 shows brain similarity scores for different mouse areas for the E// and /// conditions and the corresponding synthetic-environment conditions. There was poor correspondence between the brain similarities of the original environments and the corresponding synthetic environments. Across mouse brain areas, the correlations between the deviations from the population mean of the original and synthetic similarities were −0.473 (meadow) and −0.465 (spaceship). Interestingly, the synthetic spaceship stimuli led to high brain similarities, exceeding the scores of the original meadow stimuli (mean 0.705 vs. 0.704).

**Figure 15:**
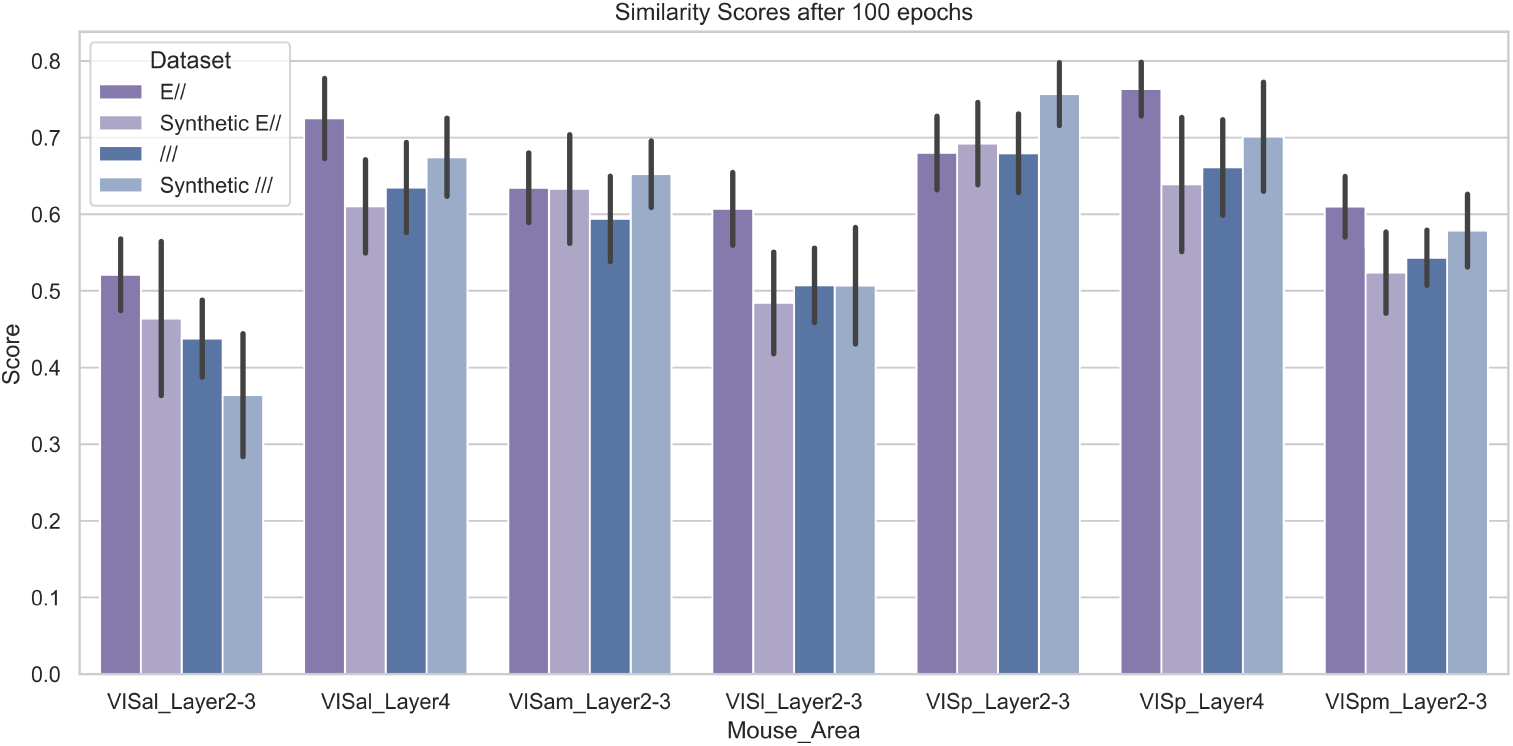
Similarity scores for the original and synthetic environments under the E// and /// conditions at end of training. Error bars show the standard deviations across random seeds.

## 4 Discussion

Deep networks are controversial (Kay, 2018; Bowers et al., 2023) models of the visual cortex, but they are unparalleled in their ability to mimic sophisticated, ethologically relevant function, and to account for stimulus-driven activity (Schrimpf et al., 2020). Critically, deep networks are not a finite set of models with fixed merits, but a space in which active exploration and innovation can lead to better alignment with the brain, and perhaps ultimately, rich mechanistic accounts of brain function. Different deep networks vary in their alignment with brain activity (Schrimpf et al., 2020), but there has been limited progress in systematically exploring the factors that affect alignment. Examples of work in this direction include Nayebi et al. (2023) and Lindsay et al. (2022a), both of which explored the impact of different training objectives.

This work experimented with three factors in stimulus statistics. We hypothesized that making certain factors in the stimuli more naturalistic would improve brain alignment, and experimented with naturalistic vs. non-naturalistic environments, motion models, and optics. The naturalistic environment consistently improved brain similarity, while the other factors had more subtle effects. Each of these three factors contributed to a statistically significant main effect and/or interaction. Across all variations, the spread in mean brain similarity was 0.062, which is over 10% of the mean similarity of untrained models.

Importantly, to confirm that changes in the simulation factors (environment, motion, and optics) impacted the model’s learning and representations, we measured the test prediction loss for each model using a combination of simulation factors different from its training condition. We consistently observed an increase in inference loss when any simulation factor differed from the training condition (Supplementary Table 4). This demonstrates that changes in any of the simulation factors altered the training data distribution. Therefore, the absence of an effect on brain similarity cannot be attributed to a lack of impact on training.

An important caveat related to motion processing is that the backbone network, MouseNet, is a two-dimensional convolutional network that processes each frame independently, and therefore cannot represent visual motion. Motion is only directly represented in the recurrent network head. It is interesting that motion statistics nonetheless impacted alignment of MouseNet with mouse brain activity. Non-naturalistic motion was dominated by visual expansion, and different static features may have provided better or worse bases for estimates of future frames in this context. However, in the mouse brain, visual motion is represented broadly, including substantial direction selectivity in VISp (Marshel et al., 2011; Niell and Scanziani, 2021). Future work should re-evaluate the impact of realistic motion statistics in a dynamic version of MouseNet.

Following common practice, we reported similarity between brain cell populations and the most similar network layers. Because there are one-to-one correspondences between mouse visual cortex cell populations and MouseNet layers, more directed comparisons are possible. However, we found that activity in a given brain cell population was not typically most aligned with the corresponding model population. This is unsurprising, in part because there is little to differentiate intermediate-level areas of MouseNet. We recently developed a version of MouseNet in which different intermediate areas cover different parts of the visual field, as they do in mouse cortex (Garrett et al., 2014). This may help to differentiate activity in these areas, and allow more meaningful comparisons between corresponding model and brain cell populations.

Our results clearly showed improved brain similarity due to training in our naturalistic environment, relative to both the artificial environment and untrained models. This raises the question of which properties of these environment might account for these results. For example, perhaps any environment with a roughly 1/f spectrum (Tolhurst et al., 1992) would yield similar results to the meadow environment. However, when we created synthetic stimuli that approximated each environment in terms of multiple such statistical properties, the resulting brain similarities were quite different than those arising from the original environments. For example, the synthetic spaceship environment led to similarities that were closer to those of the meadow environment than the spaceship environment. This suggests that the differences may be due to more complex properties of these environments.

Future work should modulate the meadow environment to explore which other properties modulate brain similarity. For example, perhaps a greater or lesser variety of features, or different density of visually distinct landmarks, would further improve brain similarity. Notably, the laboratory mice from which neural data were recorded lack experience with meadows. Perhaps training a model in a cage-like environment would yield greater similarity. Or perhaps the mouse brain is biased to perform well in more naturalistic environments, so that training a model that lacks such bias in a cage-like environment would instead worsen brain similarity.

The fact that the synthetic environments led to different brain similarities than each other also raises the possibility of optimizing brain similarity within this simplified space of visual statistics. Future work should treat the spectral, orientation, colour, and temporal autocorrelation properties as hyperparameters and optimize them for brain similarity.

Future work should continue to consider naturalistic motion and optics, due to potential interactions with other factors that we did not explore here. For example, naturalistic optics may produce different results in combination with models of extra-classical receptive field properties (Pecka et al., 2014). In general, brain similarity could be affected nonlinearly by combinations of factors. The fact that all possible interactions between factors in our study were statistically significant is an example.

Departing from much previous work that compares convolutional networks to the visual cortex, we exclusively used MouseNet (Shi et al., 2022), a model that is meant to closely approximate the feedforward structure of mouse visual cortex. In a broad comparison between models, Nayebi et al. (2023) found that MouseNet aligned more poorly with mouse brain activity than certain more conventional deep networks. However, network structure is also a factor that may interact with others to affect brain activity similarity. Thus, the factors that align models like MouseNet with the brain may be somewhat different than those that align conventional networks.

## Conclusion

We trained a deep-network model of mouse visual cortex using a self-supervised objective, and examined the impact of stimulus statistics on similarity with mouse brain activity. We used a video-game engine to create eight sets of egocentric videos as training data, and compared naturalistic vs. non-naturalistic environment, motion, and optics. Training conditions with the naturalistic environment consistently improved brain similarity relative to conditions without, and relative to untrained models. However, we could not account for these differences with simple synthetic approximations of these environments. The other factors had non-significant main effects, but significant interactions and variations across cell populations.

## Acknowledgments

Supported by the Natural Sciences and Engineering Research Council of Canada, RGPIN-05855. SB is funded by NSERC Discovery grants (RGPIN-2023-03875). Mouse motion data courtesy of Adrien Peyrache.

## Appendix

### Self-Supervised Accuracy

Table 4 shows the average self-supervised accuracy after training for 100 epochs. Each element on the diagonal is the largest element of its row/column as the evaluation dataset is the same as the training dataset.

**Table 4:** Mean self-supervised accuracy of different models (columns) when evaluated on non-matching datasets (rows)

|  |  | Model |  |  |  |  |  |  |  |
| --- | --- | --- | --- | --- | --- | --- | --- | --- | --- |
|  |  | /// | //M | /V/ | /VM | E// | E/M | EV/ | EVM |
| Evaluation Dataset | /// | 0.734182 | 0.363277 | 0.470242 | 0.231968 | 0.074951 | 0.081305 | 0.088276 | 0.084143 |
|  | //M | 0.270175 | 0.49946 | 0.262972 | 0.323558 | 0.100367 | 0.11171 | 0.11584 | 0.132935 |
|  | /V/ | 0.396715 | 0.23348 | 0.663553 | 0.292809 | 0.075493 | 0.063266 | 0.077557 | 0.082685 |
|  | /VM | 0.189918 | 0.223622 | 0.238054 | 0.460514 | 0.08571 | 0.087095 | 0.081955 | 0.106972 |
|  | E// | 0.055918 | 0.066604 | 0.081033 | 0.096017 | 0.50946 | 0.279378 | 0.305935 | 0.227143 |
|  | E/M | 0.067842 | 0.091818 | 0.112079 | 0.142089 | 0.317282 | 0.406229 | 0.235978 | 0.304871 |
|  | EV/ | 0.046551 | 0.049078 | 0.07698 | 0.085342 | 0.250058 | 0.161941 | 0.541701 | 0.228404 |
|  | EVM | 0.07332 | 0.087924 | 0.136371 | 0.131585 | 0.251107 | 0.299843 | 0.280553 | 0.414717 |

### Naturalistic motion model

Figure 16 shows part of the data we used to develop the naturalistic motion model. This is a subset of the mouse’s horizontal trajectory within a 50×50cm environment. Figures 17 and 18 compare data mouse motion data to motion patterns generated by the model.

**Figure 16:**
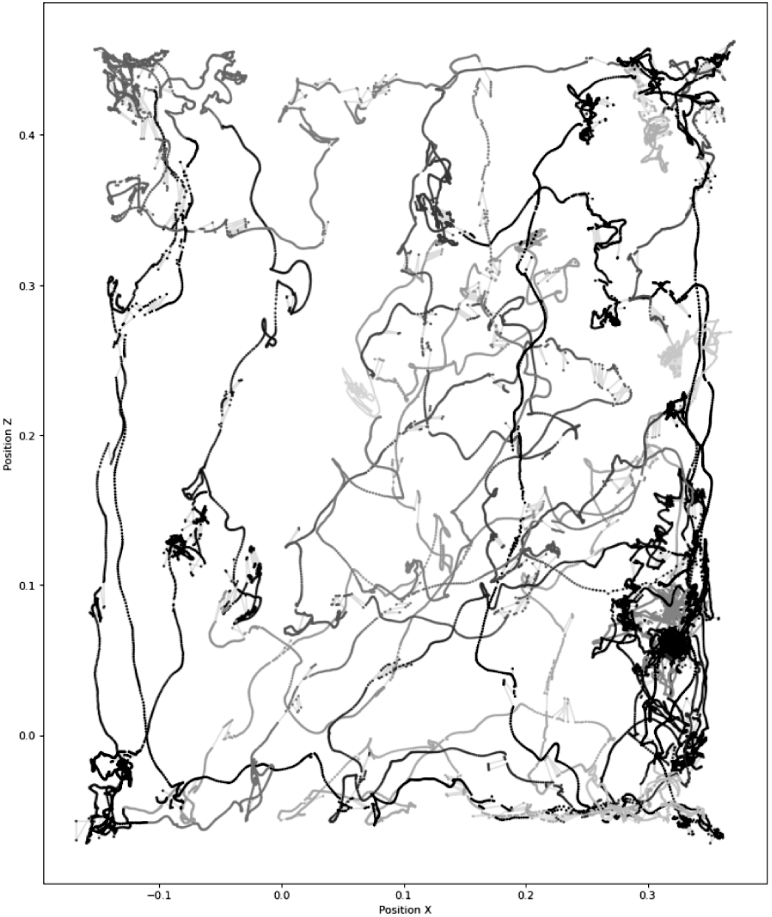
Bird’s-eye view of horizontal position from the motion capture data used to model mouse motion (courtesy of Adrien Peyrache)

**Figure 17:**
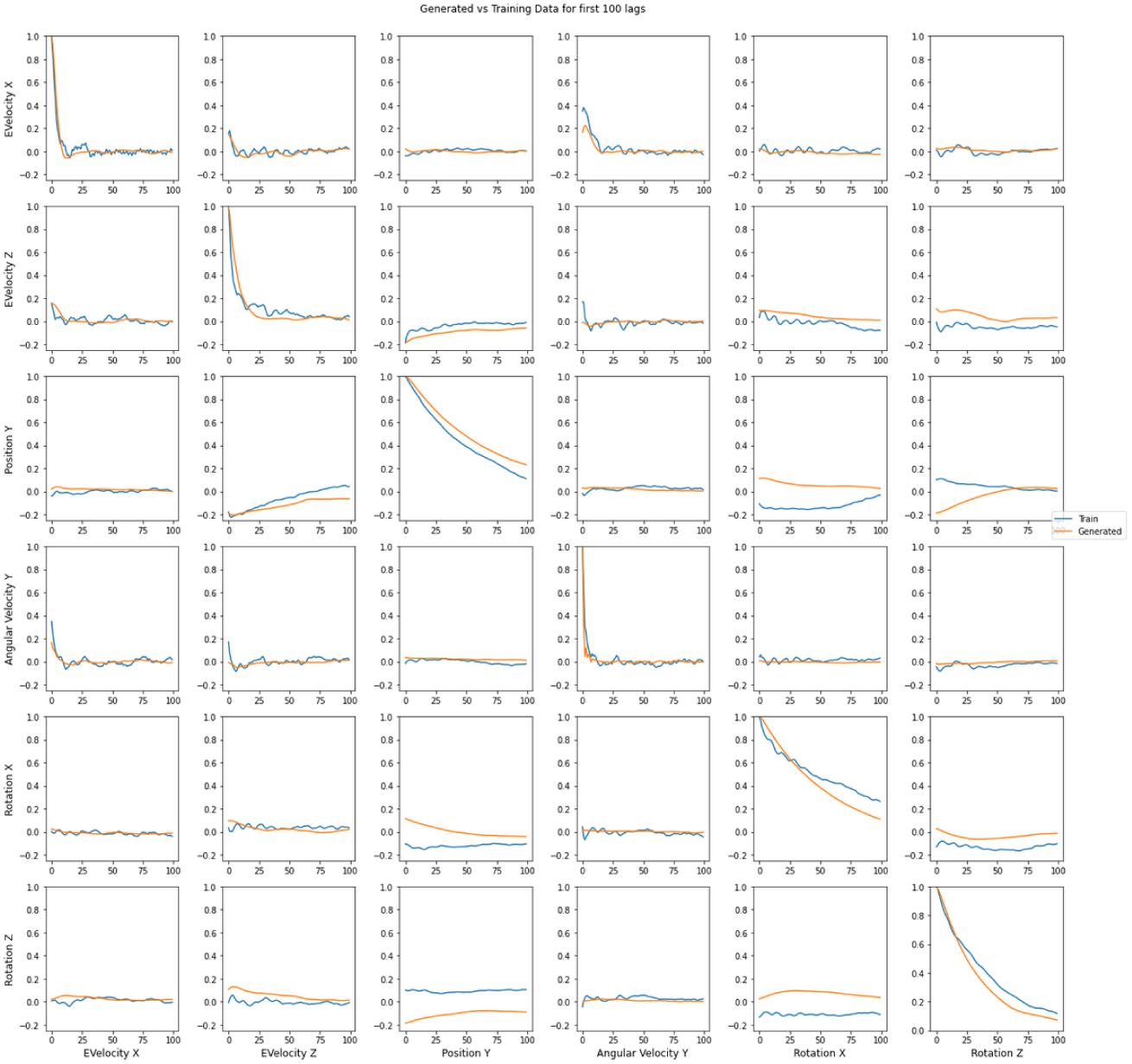
Auto and cross correlations for the first 100 lags across both the reference movement training data (blue) and the generated running data (orange)

**Figure 18:**
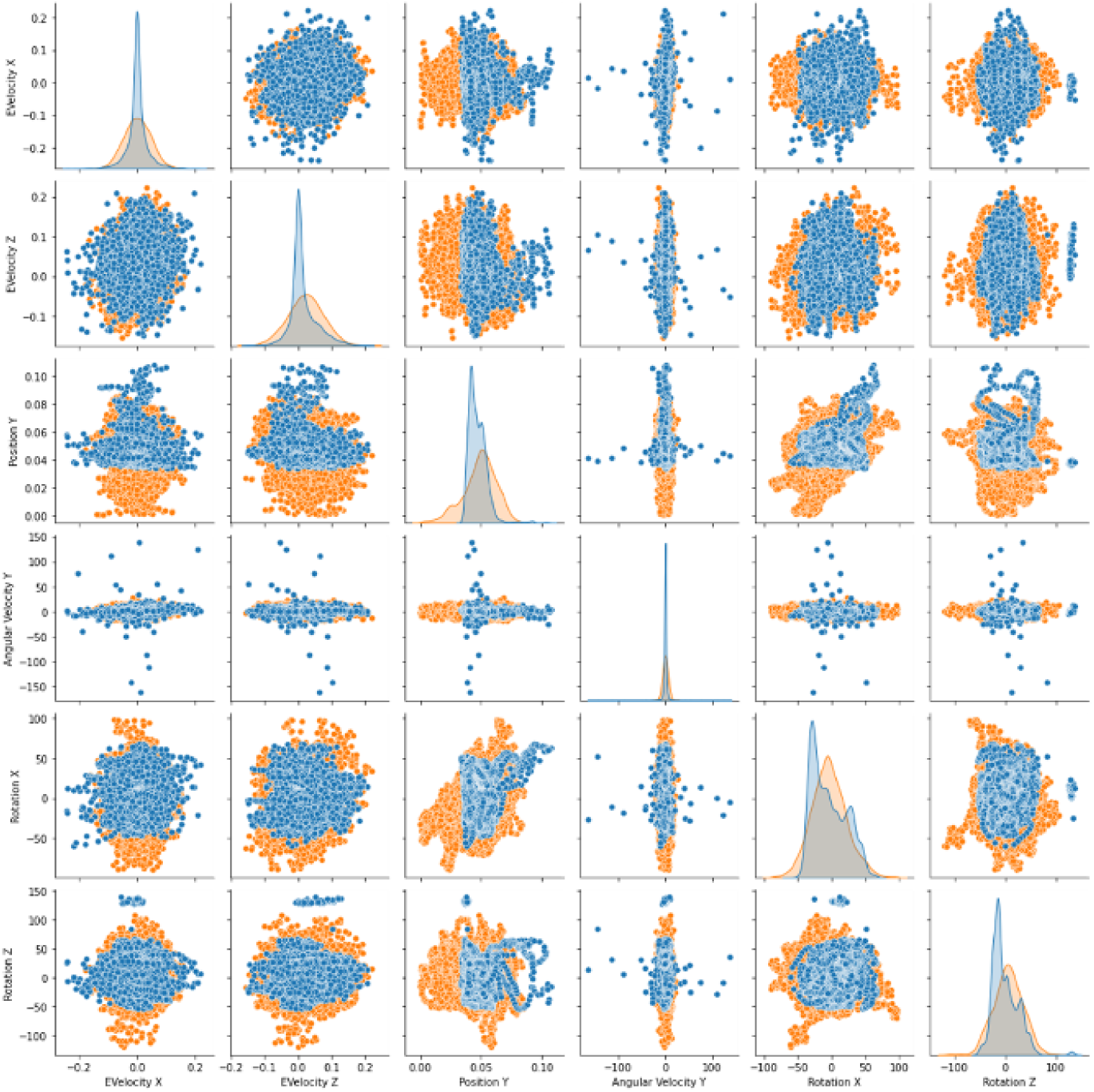
Univariate (diagonal) and bivariate (off-diagonal) distributions of motion variables in mouse data (blue) and generated running data (orange)

